# An ovine model for investigation of the microenvironment of the male mammary gland

**DOI:** 10.1101/2023.12.17.572059

**Authors:** Benjamin P. Davies, Rachael C. Crew, Anna L. K. Cochrane, Katie Davies, Andre Figueiredo Baptista, Sonja Jeckel, Ian McCrone, Youguo Niu, Benjamin W. Strugnell, Katie Waine, Abigail L. Fowden, Clare E. Bryant, John W. Wills, Dino A. Giussani, Katherine Hughes

**Affiliations:** Department of Veterinary Medicine, University of Cambridge, Cambridge, UK; Department of Physiology, Development and Neuroscience, University of Cambridge, Cambridge, UK; Department of Obstetrics and Gynaecology, University of Cambridge, UK; Farm Animal Pathology and Diagnostics, The Royal Veterinary College, Hatfield, UK; Farm Post Mortems Ltd, Durham, UK; Faculty of Veterinary Medicine, University of Calgary, Calgary, AB T3R 1J3, Canada

**Author notes:** Correspondence: Katherine Hughes, Department of Veterinary Medicine, University of Cambridge, Madingley Road, Cambridge, CB3 0ES, UK.

**Keywords:** Male, mammary gland, microenvironment, model, sheep, udder

## Abstract

The specific biology of the male breast remains relatively unexplored in spite of the increasing global prevalence of male breast cancer. Delineation of the microenvironment of the male breast is restricted by the low availability of human samples and a lack of characterisation of appropriate animal models. Unlike the mouse, the male ovine gland persists postnatally. We suggest that the male ovine mammary gland constitutes a promising adjunctive model for the male human breast. In this study we evaluate the male ovine mammary gland microenvironment, comparing intact and neutered males. Assessment of the glandular histo-anatomy highlights the resemblance of the male gland to that of the neonatal female sheep and confirms the presence of rudimentary terminal duct lobular units. Irrespective of neutered status, cell proliferation in epithelial and stromal compartments is low in males, and cell proliferation in epithelial cells and in the intralobular stroma is significantly lower than in pubertal female sheep. Between 42% and 72% of the luminal mammary epithelial cells in the male gland express the androgen receptor, and expression is significantly reduced by neutering. Luminal epithelial cells within the intact and neutered male gland also express oestrogen receptor alpha, but minimal progesterone receptor expression is observed. The distrubution of mammary leukocytes within the ducts and stroma is similar in the female mammary gland of sheep and other species. Both macrophages and T lymphocytes are intercalated in the epithelial bilayer and are more abundant in the intralobular stroma than the interlobular stroma, suggesting that they may have a protective immunological function within the vestigial glandular tissue of the male sheep. Mast cells are also observed within the stroma. These cells cluster nearer the glandular tissue and are frequently located adjacent to blood vessels. The abundance of mast cells is significantly higher in intact males compared to neutered males, suggesting that hormone signalling may impact mast cell recruitment. In this study, we demonstrate the utility of the male ovine mammary gland as a model for furthering our knowledge of postnatal male mammary biology.

## Introduction

The mammary gland nourishes and supports the development of offspring. Whilst they are intrinsically linked with female individuals, mammary glands persist in the majority of male mammals, although with notable exceptions including mice, horses and marsupials (Cardiff et al., 2018; Hughes, 2021a; Renfree et al., 1990).

Early fetal development of the male mammary gland is typically consistent with that of the female, irrespective of species (Hassiotou and Geddes, 2013; Jenkinson, 2003; Pokharel et al., 2018). Between embryonic day 11 and 13 of fetal mouse development, mammary placodes are established from ectoderm thickening on bilateral milk lines (Macias and Hinck, 2012; Stewart et al., 2019). The mammary placodes then invaginate into the mesenchyme layer, forming mammary buds (Paine and Lewis, 2017). Sexual dimorphism of the mouse mammary gland is established at embryonic day 14 (Richert et al., 2000; Stewart et al., 2019). In male mice, androgen is produced from the developing testes which causes condensation of the mesenchyme within the developing mammary gland (Dürnberger and Kratochwil, 1980; Richert et al., 2000; Vandenberg et al., 2013). This results in the morphological distortion of the mammary epithelium detaching the gland from the overlying epidermis (Drews and Drews, 1977; Richert et al., 2000; Stewart et al., 2019). The gland thereafter regresses and at birth comprises minimal vestigial glandular tissue that lacks nipples (Cardiff et al., 2018; Pokharel et al., 2018; Szabo and Vandenberg, 2021). In some genetically modified mouse strains, glandular tissue can persist after birth (Pokharel et al., 2018; Szabo and Vandenberg, 2021) but may still lack key developmental structures required for mammary gland expansion such as terminal end buds (Kolla et al., 2017).

By contrast, human prepubescent breast development is consistent between the sexes until the influence of androgen at puberty limits both ductal and stromal expansion of male breast tissue (Hassiotou and Geddes, 2013; Jesinger, 2014). The male breast comprises a small arborising ductal tree, apparently largely without well-developed lobules, embedded in an adipose-rich stroma (Fox et al., 2022).

The male breast has the potential to develop similar pathologies to that of females but at a lower incidence (Chatterji et al., 2023; Iuanow et al., 2011). However, there has been less focus on the specific biology of the male breast. Prevalence of male breast cancer is increasing globally (Fox et al., 2022) and the risk of death is significantly higher than comparable female breast cancers (Liu et al., 2018). Male breast neoplasms also have different biology compared to breast neoplasms arising in women (Chatterji et al., 2023). Research focussed specifically on the male breast is currently limited by both the low availability of human samples and the lack of characterisation of appropriate animal models in which the complete male mammary structure persists postnatally.

Sheep are commonly used to model human fetal development (Morrison et al., 2018) and we and others have noted anatomical similarities in mammary terminal duct lobular unit (TDLU) structure and stromal composition between the sheep mammary gland and the female breast (Hovey et al., 1999; Hughes, 2021b; Hughes and Watson, 2018; Nagy et al., 2021; Rowson et al., 2012). Male and female ovine fetuses exhibit parallel mammary developmental, with the gland and teat cistern present in both sexes around fetal day 80 (Jenkinson, 2003). However, there has been little examination of the male ovine mammary gland postnatally. We suggest that the mammary gland of the male sheep may constitute a promising model of the male human breast. Consequently, in this study we evaluate the mammary microenvironment in the male sheep, comparing intact and neutered males to highlight how male specific sex hormones may affect the mammary microenvironment.

## Materials and Methods

### Animals

Ovine mammary tissue was obtained from both male and female sheep that were submitted to the diagnostic veterinary anatomic pathology services of either the Department of Veterinary Medicine, University of Cambridge or to the Royal Veterinary College. Mammary tissue was also collected during the post mortem examination of Welsh mountain sheep, euthanised for research purposes under the Animals (Scientific Procedures) Act 1986. The Ethics and Welfare Committee of the Department of Veterinary Medicine, University of Cambridge, approved the study plan to use post mortem tissue in this project (references: CR223 and CR625). The nonregulated scientific use of post mortem mammary tissue collected from research animals was approved by the Named Veterinary Surgeon of the University of Cambridge. Any tissue containing evidence of mammary pathology was excluded from the study.

Male sheep used in this study ranged from 3 days old to 3 years old, but only animals older than 4 months were included in quantification analysis. Pubertal female sheep were aged from 4 months to 11 months. Mature female sheep were all older than two years and were not expected to be oestrus cycling based on the time of year of the post mortem examination (sheep are seasonal breeders). The sheep used were from a variety of breeds (Supplementary Table 1).

### Tissue processing

Mammary tissue was fixed in 10% neutral-buffered formalin for approximately one week. The entire male mammary gland and samples of female mammary parenchyma were then trimmed, processed using a routine histology protocol and embedded in paraffin. Sections were cut at 5 µm and mounted on coated glass slides (TOMO®) or stained with haematoxylin and eosin to check for microscopic pathology prior to inclusion in the study.

### Dual immunohistochemistry and immunofluorescence

Formalin fixed paraffin embedded (FFPE) tissue was subjected to antigen retrieval using a PT link module and high pH antigen retrieval solution (both Dako Pathology/Agilent Technologies, Stockport, UK). For dual immunohistochemistry, an ImmPRESS® Duet Double Staining Polymer Kit (Vector Laboratories) was used. Primary antibodies were added at the appropriate concentration (Supplementary Table 2) and incubated overnight at 4°C. Negative controls received isotype- and species-matched immunoglobulins. Counterstaining was achieved by incubating in Mayer’s Haematoxylin for 4 minutes. Slides were dehydrated in an ethanol and xylene series and Pertex® Mounting Medium was added dropwise. ClariTex Coverslips (24 x 50mm) were then applied.

For immunofluorescence, slides were first incubated with 10% normal goat serum for 1 hour at room temperature. Primary antibodies were added at the appropriate concentration (Supplementary Table 2) and incubated overnight at 4°C. Slides were then incubated in darkness with secondary antibodies (Supplementary Table 2) for 1 hour at room temperature. Negative controls received isotype- and species-matched immunoglobulins. Nuclei staining was performed by incubating with DAPI (10.9 µM) (SigmaAldrich/Merck Life Science UK Limited, Gillingham, UK) for 5 minutes. Slides were cover-slipped using Vectashield® VibranceTM Antifade mounting medium (catalogue H-1700; Vector laboratories, Peterborough, UK) and imaged using a Leica TCS SP8 confocal microscope.

### Clear, unobstructed brain imaging cocktails (CUBIC)

Mammary tissue was dissected and fixed in 10% neutral-buffered formalin for 6 to 26 hours. The gland was trimmed into smaller samples, approximately 15 x 15 x 5 mm. These samples were optically cleared using the CUBIC protocol (Lloyd-Lewis et al., 2016; Susaki et al., 2014) with the modifications outlined below. Tissue was incubated in CUBIC reagent 1A for 4 days on a shaker at 37°C. The solution was replaced daily. Samples were then incubated overnight, on at shaker at 4°C, with a blocking solution containing normal goat serum [10% (volume per volume)] and Triton X-100 [0.5% (weight per volume)] in phosphate-buffered saline (PBS). Primary antibodies, diluted in blocking solution, were applied at appropriate concentrations (Supplementary Table 2) and samples were incubated for 4 days on a shaker at 4 °C. Tissue samples were subsequently thoroughly washed in PBS and secondary antibodies (Supplementary Table 2), also diluted in blocking solution, were then applied. Samples were incubated in darkness on a shaker at 4 °C. Negative controls received isotype- and species-matched immunoglobulins. After further washing in PBS containing Triton X-100 (0.1% (weight per weight)), nuclei staining was performed by incubating in DAPI (10.9 µM) (SigmaAldrich/Merck Life Science UK Limited, Gillingham, UK) at room temperature for at least 1 hour. After further washing, CUBIC reagent 2 was applied and incubated in darkness for 4 days on shaker at 37 °C. Tissue samples were imaged in Ibidi 35 mm glass bottom dishes (catalogue 81218-200; ibidi GmbH, Gräfelfing, Germany) using a Leica TCS SP8 confocal microscope.

### Slide scanning

Slides subjected to immunohistochemical staining were scanned at 40× magnification using a NanoZoomer 2.0RS, C10730, (Hamamatsu Photonics, Hamamatsu City, Japan). Scanned sections were analysed with viewing software (NDP.view2, Hamamatsu Photonics).

### Sampling for cell proliferation and immune cell abundance

Slide scans were analysed with NDP.view2 viewing software. Depending on the tissue area available for analysis, three to eight count boxes (400 × 230 µm) were randomly placed on each slide at 1.25x magnification. Boxes containing artefacts from slide cutting or scanning were repositioned.

### Quantification of epithelial and stromal proliferation

Instances of myoepithelial and luminal cell proliferation, and proliferation of any cells located within the interlobular and intralobular stroma, as denoted by positive nuclear Ki67 staining, were manually counted. The total count for all sample boxes on the slide was calculated. This total count was then normalised to the total sampled area (mm^2^) of glandular tissue, interlobular or intralobular stroma, calculated by tracing around the area using the NDP.view2 freehand annotation tool. The protocol was repeated for all individuals within each experimental group and the mean was derived.

### Manual quantification of hormone receptor expression

A Leica TCS SP8 confocal microscope was utilised to produce x400 magnification tile scans of immunofluorescence slides. Tile scans were analysed using Fiji (Schindelin et al., 2012), and, depending on the size of the tissue, three to eight count boxes (each measuring 400 × 230 µm) were randomly placed on each slide at low magnification. Hormone receptor positive luminal epithelial cell nuclei were manually counted and the total count for all sample boxes on the slide was calculated. This count was expressed as a percentage of the total number of luminal epithelial cell nuclei within all sample count boxes. The protocol was repeated for all individuals within each experimental group and the mean percentage of hormone receptor positive luminal cell nuclei for each group was calculated.

### Epithelial and Stromal Macrophage and T lymphocyte abundance

Using NDP.view2, the number of epithelial-associated IBA1 positive-macrophages and CD3-positive T lymphocytes were counted. Cells were considered epithelial-associated if >50% of their cytoplasmic perimeter contacted the basement membrane. The total number of epithelial-associated macrophages and T lymphocytes in all count boxes was calculated for each individual, together with the total number of epithelial cells, and the epithelial-associated immune cell count was expressed per 100 luminal epithelial cells. Stromal macrophages and T lymphocytes were considered to be interlobular and intralobular respectively when >50% of their cytoplasmic perimeter was within the respective type of stroma. The total number of macrophages and T lymphocytes for each stroma type was calculated. This count was normalised to the total sampled area (mm^2^) of interlobular or intralobular stroma, determined using the NDP.view2 freehand annotation tool. The protocol was repeated for all individuals within each experimental group and the mean was derived.

### Statistical analysis

To assess statistical significance in comparisons between cell proliferation and comparisons between hormone receptor expression, a Kruskal-Wallis test was conducted. Statistical significance in the differences in immune cell abundance between intact and neutered males were assessed using a Mann-Whitney U test. To assess statistical significance in the differences in immune cell abundance between intra- and interlobular stroma, a Wilcoxon signed-rank test was performed. All data were collected using Microsoft Excel and were analysed using R studio (RStudio: Integrated Development for R. RStudio, PBC, Boston, MA URL http://www.rstudio.com/). All data are presented as mean values + the standard deviation.

## Results and Discussion

### The gross and histo-anatomy of the male sheep mammary gland resembles that of the neonatal female

In total, thirty-three sheep were analysed in the study (Suppl. Table 1) Nineteen of these sheep were male, comprising eight intact males and eleven neutered males. These male sheep ranged between three days old to three years old, but only male sheep older than four months were included in analyses involving quantification. These male sheep were of several different breeds, including four Cheviot cross males, three Welsh mountain males, three Texel cross males and one of each of Jacob cross, New Zealand Romney, Suffolk, Mule, Mule cross and Shetland. For three male sheep no breed information was available. Male sheep were compared to a group of seven pubertal female sheep and seven mature female ewes. Pubertal females ranged from four to eleven months in age. These ewes were of three different breeds comprising five Welsh mountain ewes and one each of Texel cross and Beltex. Mature females were all older than two years and all seven were Welsh mountain ewes.

To investigate the possibility that the mammary gland of the male sheep is a useful adjunct model for the male breast, we initially characterised the gross and histo-anatomy of the gland. The male sheep mammary gland is located in the sub-epithelial tissues below the teat (Fig. 1 a) and is composed of branching ducts that terminate in terminal duct lobular units (TDLUs) (Fig. 1 b-e). The walls of the ducts comprise a bilayer of luminal epithelial cells and basal myoepithelial cells (Fig. 1 b). Both intra- and interlobular mammary stroma surrounds the ductal system (Fig. 1 c) and, like that of the female ruminant, expresses smooth muscle actin (Nagy et al., 2021; Safayi et al., 2012). Overall the structure of the male gland is very similar to that of the neonatal female sheep (Hughes, 2021b; Nagy et al., 2021) and contrasts the minimal vestigial glandular tissue present in the postnatal male mouse mammary gland (Cardiff et al., 2018; Pokharel et al., 2018; Stewart et al., 2019; Szabo and Vandenberg, 2021).

**Figure 1:**
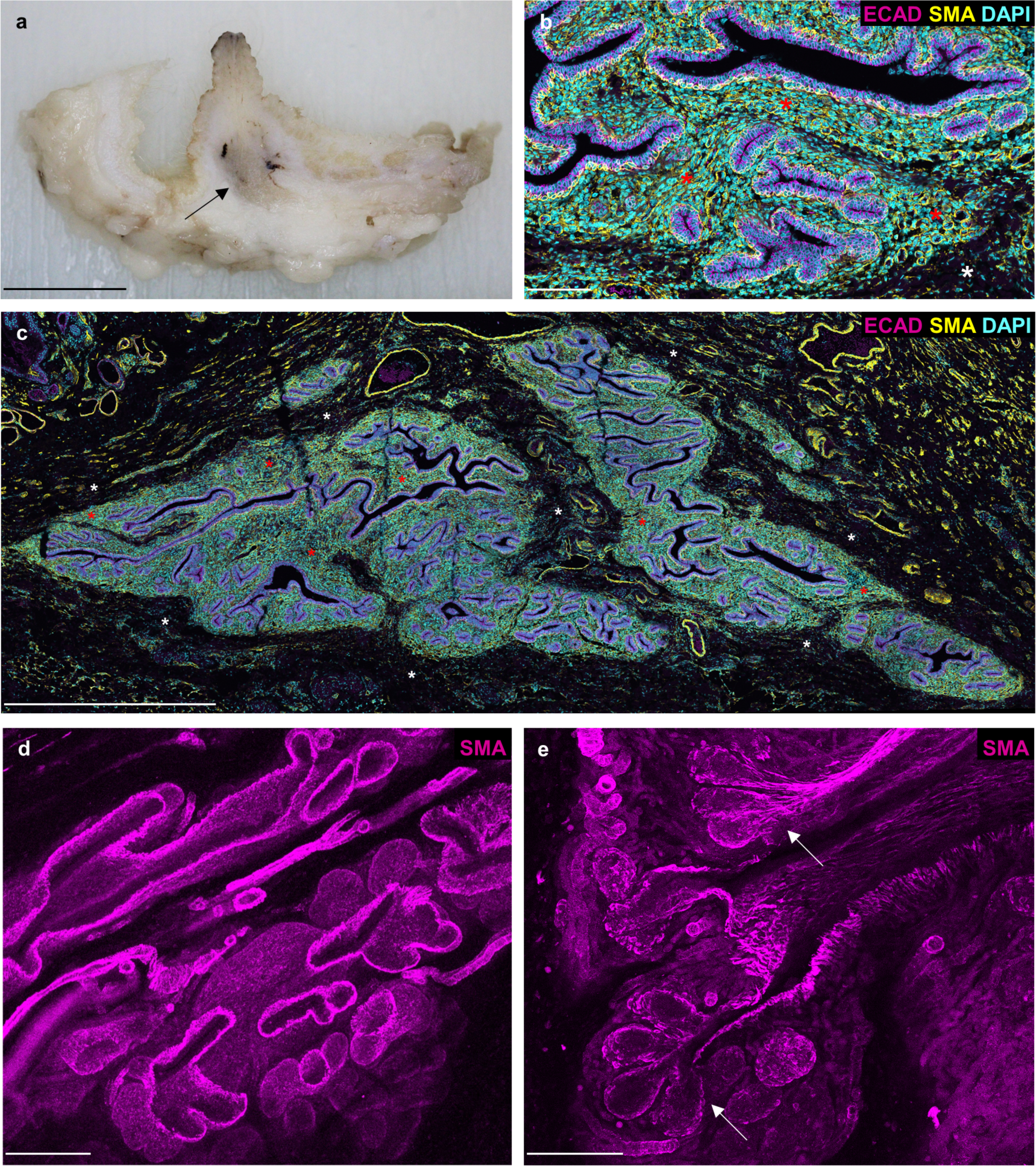
Macro and histo-anatomy of the male ovine mammary gland. (**a**) A sub-gross image of fixed male ovine mammary tissue. Arrow indicates mammary gland. (**b**, **c**) Immunofluorescence staining for luminal epithelial cells, using E-cadherin (magenta), myoepithelial cells, using alpha-smooth muscle actin (yellow) and DNA, using DAPI (cyan). Red asterisks indicate areas of intralobular mammary stroma. White asterisks indicate areas of interlobular mammary stroma. (**d**, **e**) 3D maximum intensity projections of optically cleared male ovine mammary tissue, using confocal microscopy. Immunofluorescence staining for alpha-smooth muscle actin (magenta). Arrows indicate terminal duct lobular units. Images are representative of 3 biological repeats Scale bar = 1 cm (**a**); 100 µm (**b**); 1 mm (**c**); 200 µm (**d**); 100 µm (**e**).

### The male ovine mammary gland exhibits minimal cell proliferation, irrespective of neutered status

Given that the female mammary gland undergoes dynamic changes in cell proliferation throughout its postnatal development cycle (Inman et al., 2015) we wished to assess the proliferation dynamics of different cellular compartments within the male gland, comparing intact and neutered male animals with pubertal females and mature females. Proliferation in both epithelial and stromal compartments is similarly low in male sheep irrespective of neuter status. Proliferation in luminal epithelial cells, myoepithelial cells and cells present in the intralobular stroma, is significantly higher in pubertal females when compared to both intact and neutered males (Fig 2 e-g).

**Figure 2:**
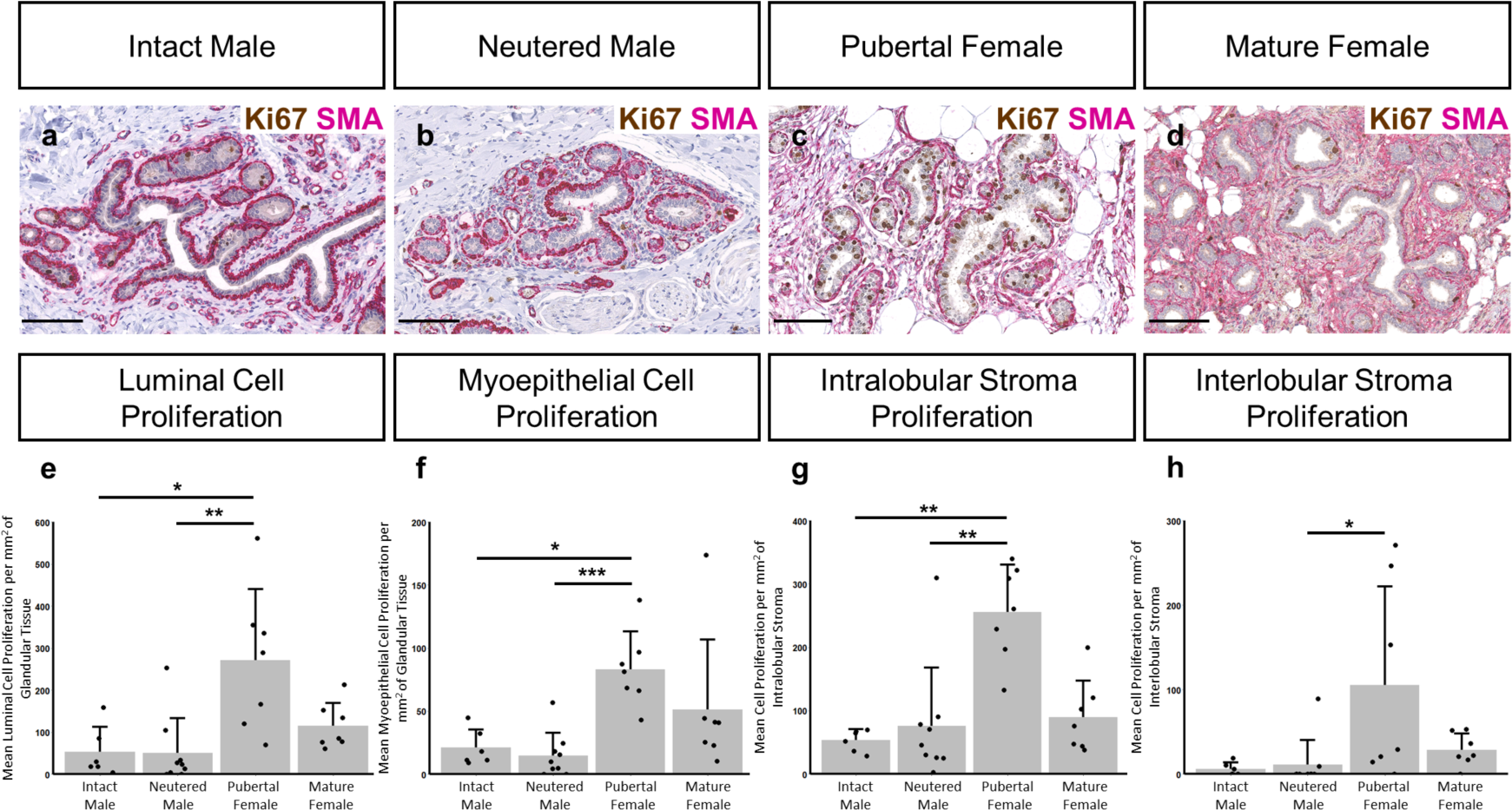
Proliferation dynamics within the intact male, neutered male, pubertal female and mature female mammary gland. (**a-d**) Dual immunohistochemical staining for Ki67 (brown) and alpha-SMA (magenta) in the mammary gland of intact males (**a**), neutered males (**b**), pubertal females (**c**) and mature females (**d**). (**e-h**) Bar graphs illustrating differences in mean luminal cell proliferation (**e**), myoepithelial cell proliferation (**f**), intralobular stroma proliferation (**g**) and interlobular stroma proliferation (**h**) per mm^2^ of glandular tissue, intra- or interlobular mammary stroma + standard deviation (*p< 0.05, **p< 0.01, ***p< 0.001, N= 6 for intact males, N =9 for neutered males, N= 7 for pubertal females, N = 7 for mature females, using Kruskal-Wallis test). Dots represent individual sheep. Images representative of 6 (**a**), 9 (**b**) and 7 (**c, d**) biological repeats. All IHC shown with a haematoxylin counterstain. Scale bar = 100 µm (**a-d**).

### Neutering decreases mammary androgen receptor expression in sheep

In our study population, between 42% and 72% of luminal mammary epithelial cells in intact male sheep express androgen receptor (AR) and expression is abrogated by neutering (Fig. 3). Previously studies have illustrated that androgen receptor activation limits the expansion of glandular tissue within the female mammary gland. Transgenic mouse models in which the production or action of AR was ablated, exhibited increased cell proliferation, ductal branching and number of terminal end buds (Gao et al., 2014; Simanainen et al., 2012). In humans, AR functions to limit the growth of breast tissue in both prepubescent boys and girls (Dimitrakakis and Bondy, 2009), and our data in the sheep are consistent with this observation.

**Figure 3:**
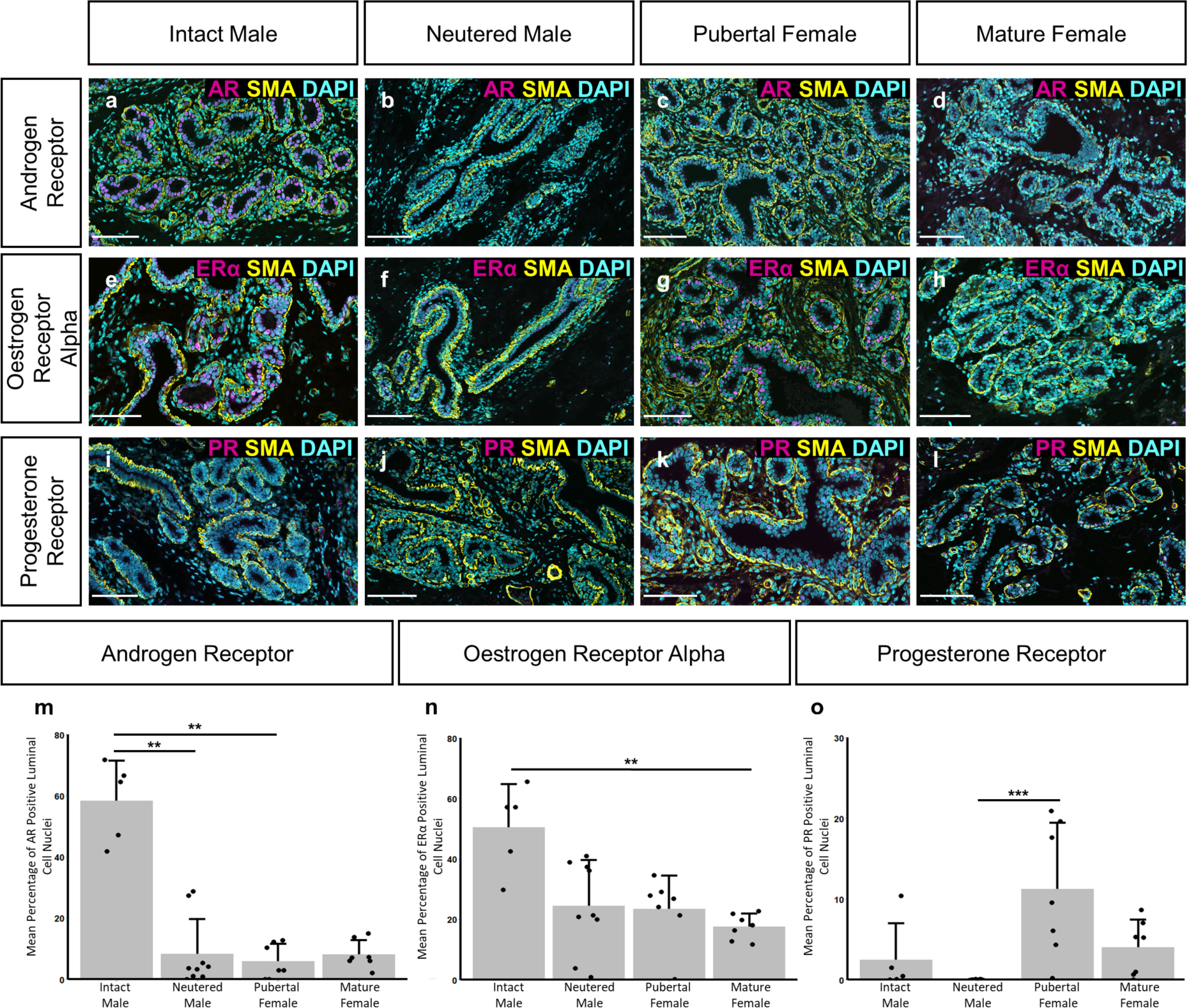
Differences in mammary epithelial hormone receptor expression between intact males, neutered males, pubertal females and mature females. **(a-l)** Immunofluorescence staining for androgen receptor (AR) **(a-d),** oestrogen receptor alpha (ERα) **(e-h)**, progesterone receptor A/B (PR) **(i-l)** (magenta), alpha-SMA (yellow) and DAPI (cyan) in the mammary gland of intact males (**a, e, i**), neutered males (**b, f, j**), pubertal females (**c, g, k**) and mature females (**d, h, l**). (**m-o**) Bar graphs illustrating differences in the mean percentage of AR (**m**), ERα (**n**) or PR (**o**) positive luminal cell nuclei + standard deviation (**p< 0.01, ***p< 0.001, N= 5 for intact males, N =9 for neutered males, N= 7 for pubertal females, N = 7 for mature females, using Kruskal-Wallis test). Dots represent individual sheep. Images representative of 5 (**a, e, i**), 9 (**b, f, j**) and 7 (**c, d, g, h, k, l**) biological repeats. Scale bar = 100 µm (**a-l**).

Oestrogen receptor alpha (ER alpha) expression has previously been highlighted in epithelial cells within the TDLUs of prepubertal sheep (Colitti and Parillo, 2013) and is similarly observed in both male and female groups in our study (Fig. 3e-3h). Our analysis indicates that intact males have a higher percentage of luminal epithelial cells exhibiting ER alpha expression than mature females (Fig. 3n), potentially reflecting a variable composition of luminal epithelial sub-groups in the male gland compared to the female. Understanding the balance between AR and ER alpha expression in the male mammary gland is important because the vast majority of male breast cancer is ER positive (Cardoso et al., 2018; Chatterji et al., 2023).

Mean progesterone receptor expression is higher in pubertal females than neutered males (Fig. 3o). Progesterone receptor expression has been previously reported in the alveolar cells of the female ovine mammary gland during lactation (Colitti and Parillo, 2013) and prior research, using hormone treated murine mammary glands, indicates that progesterone receptor signalling promotes ductal branching during puberty (Atwood et al., 2000).

Progesterone receptor signalling has also been shown to promote cell proliferation in multiple mammary cell types during puberty and pregnancy (Atwood et al., 2000; Brisken et al., 1998; Hilton et al., 2015). The low expression of progesterone receptor in neutered males and low but variable levels of progesterone receptor expression in intact males are consistent with the relatively limited tissue area occupied by the male gland and the lack of cell proliferation we observe within the male mammary gland (Fig. 2e-2h).

### Macrophages and T lymphocytes are intercalated in the epithelial bilayer and cluster nearer ductal structures in the male ovine mammary gland

In both intact and neutered male sheep, IBA-1 positive macrophages are intercalated in the epithelial bilayer and are present in the intra- and interlobular mammary stroma. There are abundant macrophages surrounding the TDLUs and many are intimately associated with the epithelium (Fig. 4). The localisation of mammary macrophages in the male sheep is overall consistent with previous characterisation of the female neonatal and pubertal ovine mammary gland, with the exception that the macrophages within the male gland do not appear to exhibit the periodicity previously noted in females (Nagy et al., 2021). The intraepithelial macrophages of the male gland may be similar to the ductal macrophages reported in the mouse mammary gland (Dawson et al., 2020). These ductal macrophages have phenotypes and gene expression patterns distinct from stromal macrophages, reflecting an immune surveillance function (Dawson et al., 2020). The macrophages within the male ovine gland could have a similar role, but their precise phenotype and function requires further elucidation. In female mice, macrophage abundance also is affected by oestrogen and progesterone, during oestrus cycling (Chua et al., 2010; Hodson et al., 2013; Tower et al., 2022) and macrophages are receptive to androgen signalling (Liva and Voskuhl, 2001). However, our data suggest that male ovine mammary gland macrophage abundance is unaffected by neutered status (Suppl. Fig 1), an observation warranting further future investigation.

**Figure 4:**
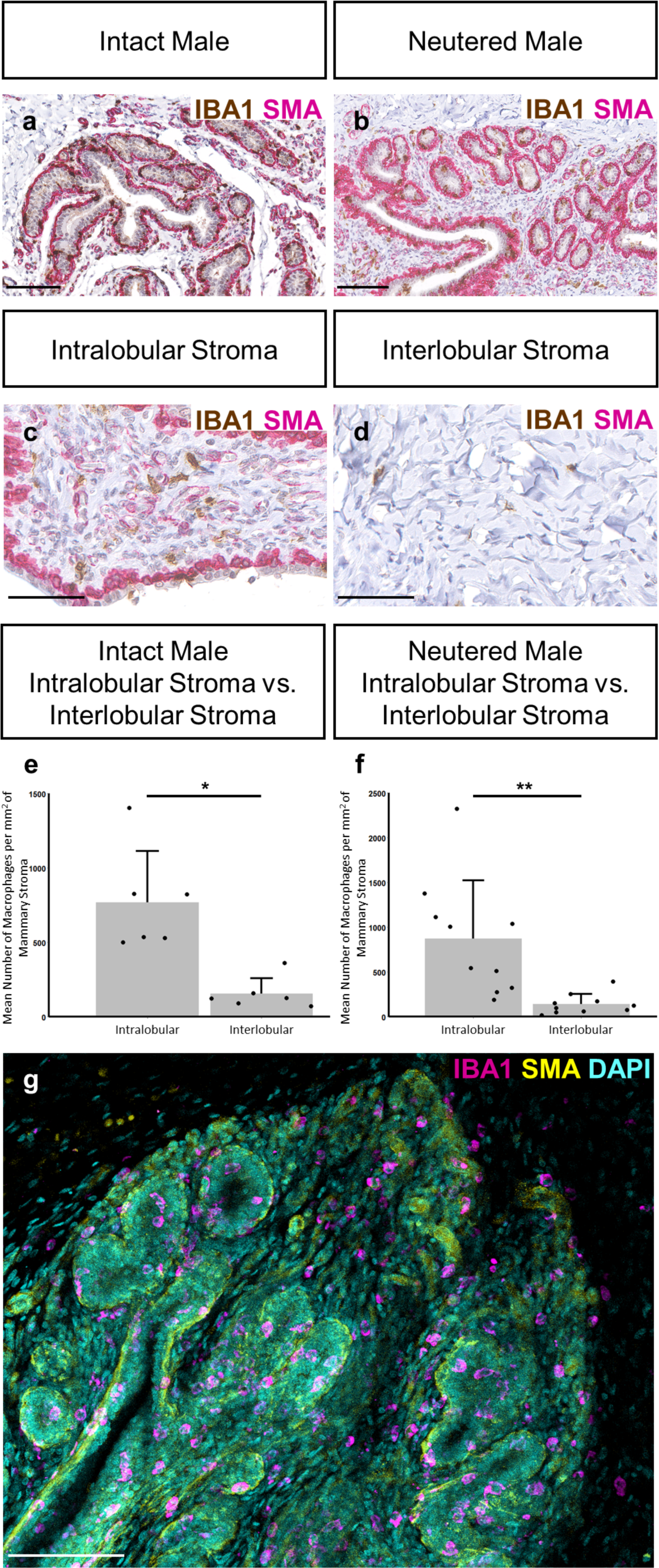
Macrophage abundance and localisation within the intact and neutered male ovine mammary gland. (**a-d**) Dual immunohistochemical staining for IBA1 (brown) and alpha-SMA (magenta) in the mammary gland of intact males (**a**) and neutered males (**b**). (**c-d**) The abundance of macrophages in the intralobular (**c**) and interlobular mammary stroma (**d**). (**e, f**) Bar graphs illustrating differences in the mean number of macrophages per mm^2^ of mammary stroma in the intralobular and interlobular stroma of intact (**e**) and neutered males (**f**) + standard deviation (*p< 0.05, **p< 0.01, N= 6 for intact males, N =10 for neutered males, using Wilcoxon signed-rank test). Dots represent individual sheep. (**g**) A 3D maximum intensity projection of optically cleared male ovine mammary tissue, using confocal microscopy. Immunofluorescence staining for IBA1 (magenta), alpha-SMA (yellow) and DAPI (cyan). Images representative of 6 (**a**), 10 (**b, c, d**) and 3 (**g**) biological repeats. Scale bar = 100 µm (**a, b, g**); 50 µm (**c, d**).

In both intact and neutered males, the abundance of macrophages is significantly higher in the intralobular stroma, directly surrounding the glandular tissue, compared to the more distant interlobular stroma (Fig. 4). This is consistent with previous examination of the female bovine mammary gland (Beaudry et al., 2016).

Similar to epithelial-associated macrophages, CD3 positive T lymphocytes are intercalated in the epithelial bilayer and are present in the intra- and interlobular mammary stroma. In contrast, there are few CD20-positive B lymphocytes (Fig. 5). Comparisons between intact and neutered males highlight that there is no significant difference in abundance of T lymphocytes associated with the epithelium or in the intra- or interlobular stroma (Suppl. Fig. 2). In neutered males there are more T lymphocytes in the intralobular stroma than in the interlobular stroma (Fig. 5). The clustering of both macrophages and T lymphocytes within and near the male mammary ductal structures may imply that they have an active immunological function with the male ovine vestigial glandular tissue, which has a potential portal for ingress of pathogens from the exterior through the teat.

**Figure 5:**
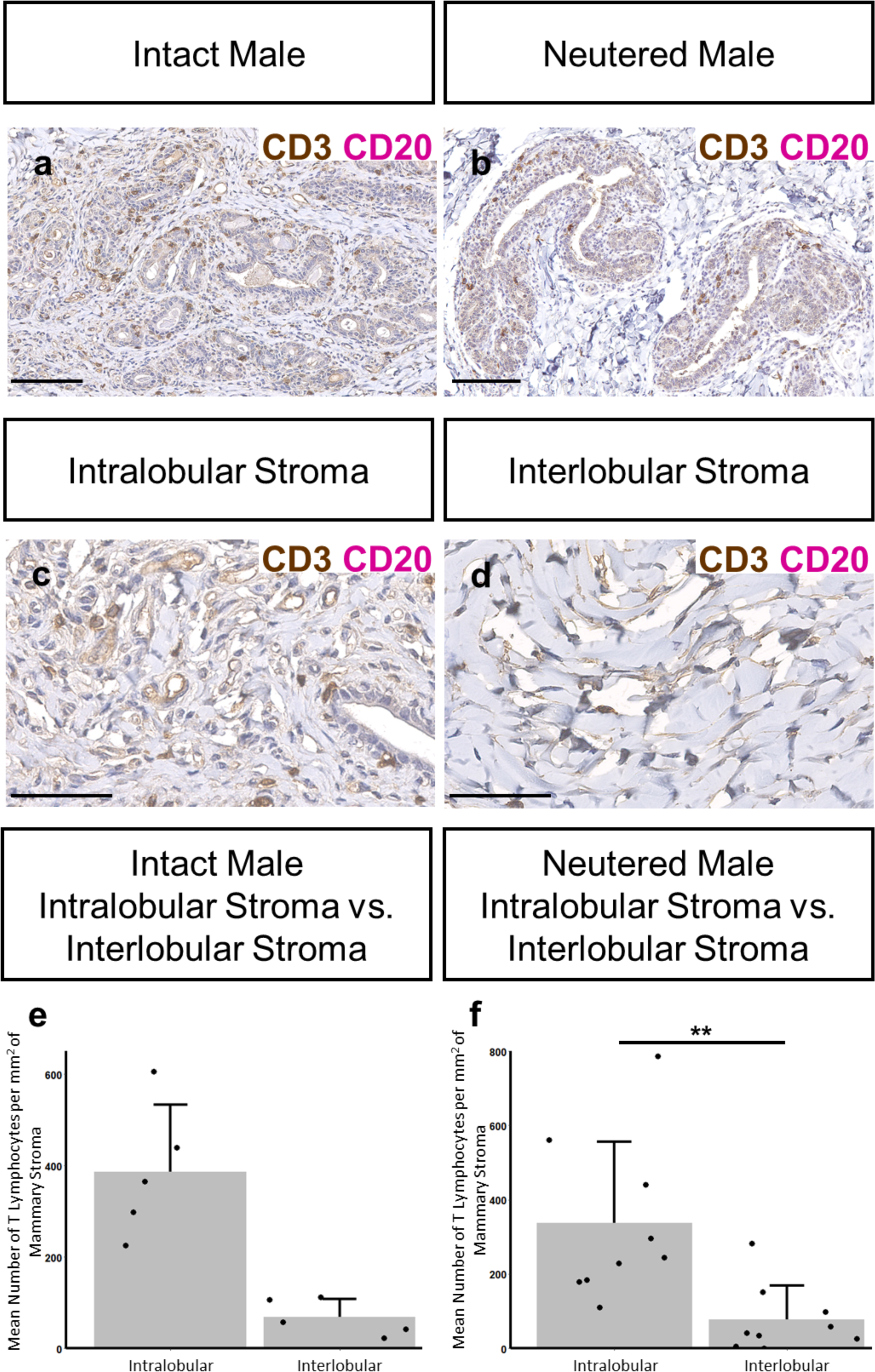
T-lymphocyte abundance within the intact and neutered male ovine mammary gland. (**a-d**) Dual immunohistochemical staining for CD3 (brown) and CD20 (magenta) in the mammary gland of intact males (**a**) and neutered males (**b**) and within the intralobular (**c**) and interlobular mammary stroma (**d**). (**e, f**) Bar graphs illustrating differences in the mean number of T-lymphocytes per mm^2^ of mammary stroma in the intralobular and interlobular stroma of intact (**e**) and neutered males (**f**) + standard deviation (**p< 0.01, N= 5 for intact males, N =9 for neutered males, using Wilcoxon signed-rank test). Dots represent individual sheep. Images representative of 5 (**a, c, d**) and 9 (**b**) biological repeats. All IHC shown with a haematoxylin counterstain. Scale bar = 100 µm (**a, b**); 50 µm (**c, d**).

### Mast cells cluster nearer ductal structures in the male ovine mammary gland and their abundance is significantly higher in intact males

Mast cells have been previously identified in the mammary gland of the mouse (Hughes et al., 2012; Lilla and Werb, 2010), rat (Ramirez et al., 2012) and cow (Beaudry et al., 2016). Typically, histological identification of mast cells is carried out using toluidine blue staining to highlight the cells’ metachromatic staining granules. We and others have used toluidine blue to identify mast cells in the mammary gland of laboratory rodents (Hughes et al., 2012; Lilla and Werb, 2010; Ramirez et al., 2012). However, toluidine blue staining does not positively stain mast cells in the ovine mammary gland, potentially due to differences in granule composition (Suppl. Fig. 3). Consequently, we have stained for c-Kit, a transmembrane tyrosine kinase receptor (Ribatti, 2018), that has been previously used to identify mast cells in human tissues (Lammie et al., 1994). Mast cells are present in the mammary stroma in both intact and neutered male glands and are present in close proximity to blood vessels (Fig. 6). This localisation is consistent with previous descriptions of mast cells in the female mouse mammary gland (Lilla and Werb, 2010).

**Figure 6:**
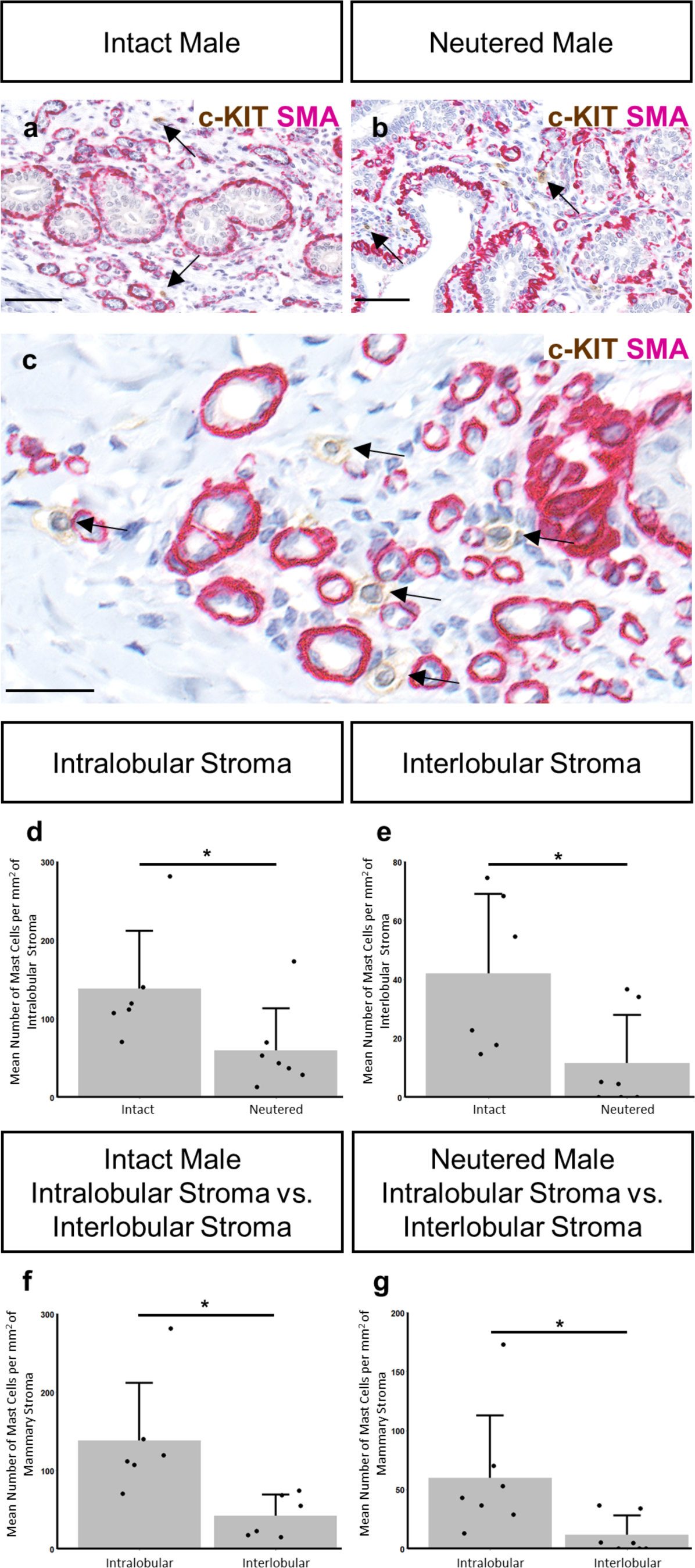
Mast cells abundance is significantly higher in intact males. (**a-c**) Dual immunohistochemical staining for c-Kit (brown) and alpha-SMA (magenta) in the mammary gland of intact males (**a**) and neutered males (**b**). Arrows indicate positive staining for mast cells. (**c**) Mast cells are located in close proximity to blood vessels in the male ovine mammary gland. Arrows indicate positive mast cell staining. (**d, e**) Bar graphs illustrating differences in the mean number of mast cells per mm^2^ of intralobular (**d**) and interlobular (**e**) stroma in intact and neutered males + standard deviation (*p< 0.05, N= 6 for intact males, N =7 for neutered males, using Mann-Whitney U test). (**f, g**) Bar graphs illustrating differences in the mean number of mast cells per mm^2^ of mammary stroma in the intralobular and interlobular stroma of intact (**f**) and neutered males (**g**) + standard deviation (*p< 0.05, N= 6 for intact males, N =7 for neutered males, using Wilcoxon signed-rank test). Dots represent individual sheep. Images representative of 6 (**a, c**) and 7 (**b**) biological repeats. All IHC shown with a haematoxylin counterstain. Scale bar = 50 µm (**a, b**); 25 µm (**c**).

The abundance of mast cells in the intra- and interlobular stroma is significantly higher in intact males compared to neutered males (Fig. 6d - e), suggesting that steroid hormone signalling may impact mast cell recruitment. A prior study highlighted that mast cells within the human skin express androgen receptors, but mast cell degranulation remained unchanged upon the administration of a testosterone treatment (Chen et al., 2010). However, authors did not comment how testosterone treatment may have affected mast cell abundance. There is also some evidence that oestrogen could affect mast cell recruitment in the female bovine mammary gland, where researchers identified a trend in which that mast cell number increases upon the exogenous oestrogen treatment (Beaudry et al., 2016). Elucidating how specific hormone receptor signalling may affect mast cell recruitment within the male ovine gland is a direction for further experimental investigation.

Consistent with the analysis of macrophage and T lymphocyte abundance, mast cell abundance is also significantly higher in the intralobular stroma, compared to the more distant interlobular stroma (Fig. 6f - g). This is seen in both intact and neutered males. Similarly in the rat mammary gland mast cells tend to be located in the stroma surrounding the ducts (Ramirez et al., 2012). Mast cell number is also four times higher in the stroma adjacent to the mammary ducts, compared to more distant regions, in the female bovine mammary gland (Beaudry et al., 2016). This suggests that mammary mast cell localisation is similar between species and between males and females.

### Conclusions

The ovine tissue analysed in this study was obtained from multiple sources from sheep with different genetic backgrounds and maintained with different husbandry practices. Additionally, tissue obtained from pubertal females was collected from sheep euthanised throughout the year and their stage of the oestrus cycle at the time of tissue collection is unknown. Together, this constitutes a heterogeneous sample population which introduces considerable variability into the dataset. Arguably, this mirrors the heterogeneity in the mammary microenvironment in the human population.

This study demonstrates the utility of the male ovine mammary gland as a tool to further our understanding of postnatal male mammary biology. This vestigial glandular structure contains a diverse set of immune cell types and exhibits distinct hormone receptor expression patterns, features that in both cases are affected by neutered status. Interestingly, the immune microenvironment of the male ovine gland appears to share features with that of the female gland in sheep and other species. This observation, together with the well-documented histo-anatomical similarities of the sheep mammary gland to the human breast (Hughes, 2021b; Nagy et al., 2021; Rowson et al., 2012) indicate that the male sheep mammary gland may be a useful adjunctive model of the male human breast. As mammary tumourigenesis is rare in sheep (Hughes, 2021b; Newman et al., 2021), the male ovine gland does not offer a direct model for male breast cancer, but it will facilitate furthering understanding of normal male mammary biology to use in comparative studies. In addition, the relative resistance of the sheep to development of mammary tumours may be a further fruitful avenue of future comparative study (Hughes, 2023).

## Acknowledgements

BPD is supported by an Anatomical Society PhD studentship awarded to KH. The authors would like to thank Hugh Balmer, Yvonne Pratt, Andrea Starling and Emma Ward of the Department of Veterinary Medicine, University of Cambridge, for their excellent technical expertise in the preparation of tissue sections, and Mathew Rhodes, of the same department, for technical assistance in the post mortem room. We acknowledge the Ethics and Welfare Committee of the Department of Veterinary Medicine, University of Cambridge, for their review of the study plan relating to the use of tissue from ovine post mortem examinations for the study of mammary gland biology (references: CR223 and CR625). Parts of this data were presented in oral abstract form at a the European Network for Breast Development and Cancer online seminar series (presentation date 6 December 2022) and at the Anatomical Society Summer Meeting 2023 in Bangor, Wales (25-27 July 2023; presentation date 25 July 2023).

## Author contributions

BPD and KH contributed to study concept and design. BPD, RCC, ALKC, KD, AFB, SJ, IM, YN, BWS, KW, ALF, DAG and KH contributed to acquisition of data. BPD, CEB, JWW, and KH contributed to data analysis and interpretation. BPD and KH drafted the manuscript. All authors contributed to critical revision of the manuscript and approved the final version.

## Supplementary Material

**Supplementary Figure 1:**
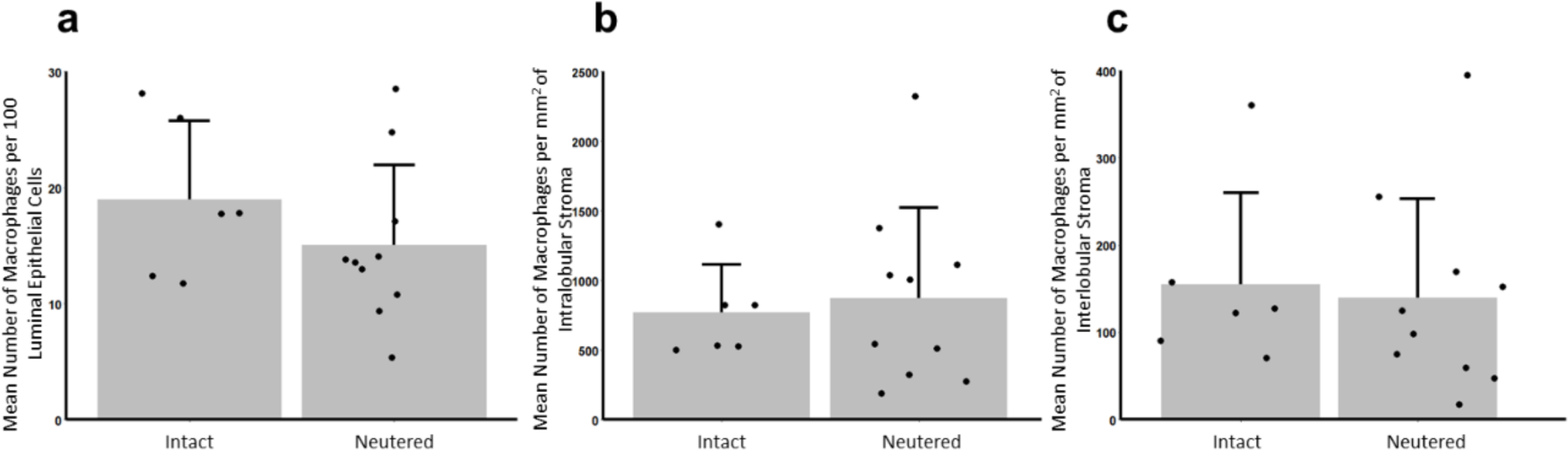
There is no significant difference in macrophage abundance in the mammary glands of intact and neutered males. (a-c) Bar graphs illustrating differences, between intact and neutered males, in the mean number of macrophages per 100 luminal epithelial cells (a) and the mean number of macrophages per mm^2^ of intralobular (b) or interlobular (c) + standard deviation (N= 6 for intact males, N =10 for neutered males, using Mann-Whitney U test). Dots represent individual sheep.

**Supplementary Figure 2:**
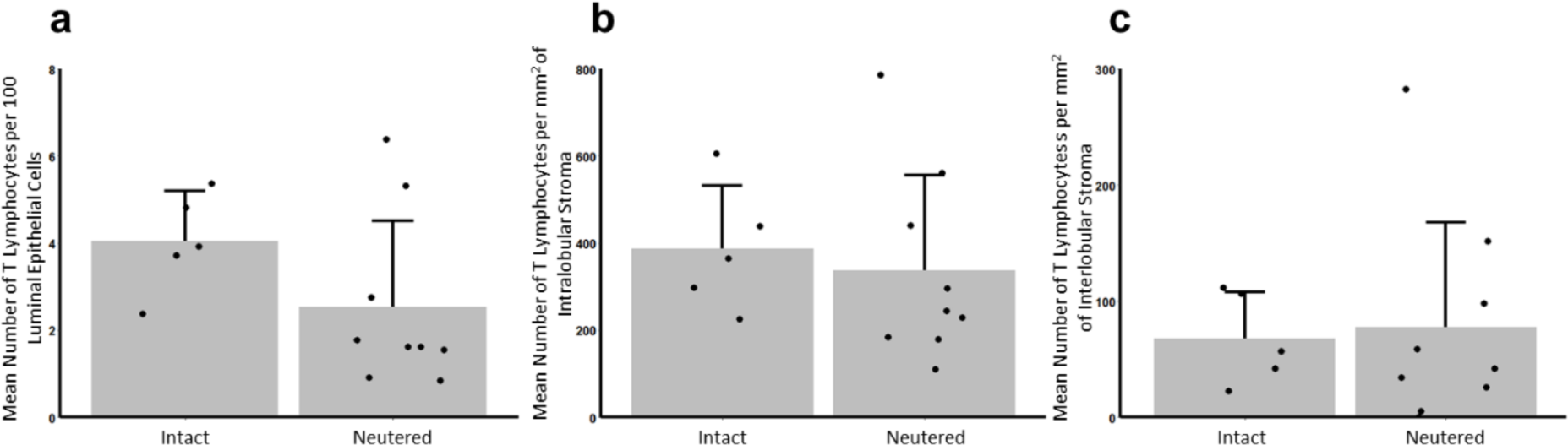
There is no significant difference in T-lymphocyte abundance in the mammary glands of intact and neutered males. (a-c) Bar graphs illustrating differences, between intact and neutered males, in the mean number of T-lymphocytes per 100 luminal epithelial cells (a) and the mean number of T-lymphocytes per mm^2^ of intralobular (b) or interlobular (c) + standard deviation (N= 5 for intact males, N =9 for neutered males, using Mann-Whitney U test). Dots represent individual sheep.

**Supplementary Figure 3:**
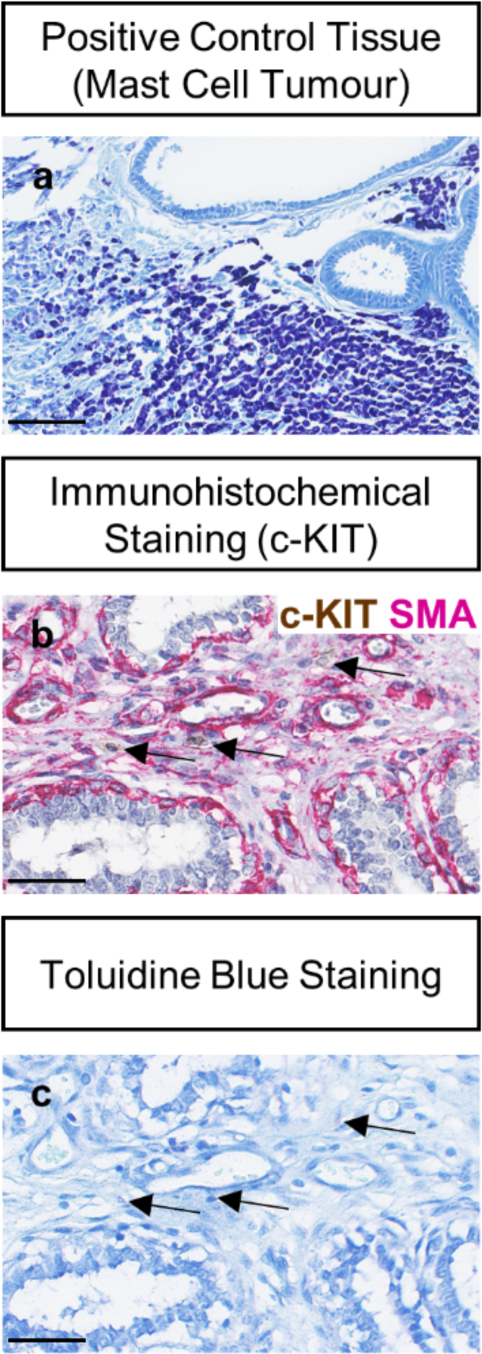
Toluidine blue staining does not positively stain mast cells in the ovine mammary gland. (a) Toluidine blue staining in a positive control tissue, a mast cell tumour. (b, c) Sequentially cut sections of FFPE ovine mammary gland tissue. (b) Dual immunohistochemical staining for c-Kit and alpha-SMA. Arrows indicate positive staining for mast cells. (c) Toluidine blue staining. Arrows indicate areas where positive toluidine staining should be. Images representative of 3 biological repeats. All IHC shown with a haematoxylin counterstain. Scale bar = 100 µm (a); 50 µm (b, c).

**Supplementary Table 1:** Full details of sheep used in the study.

| Sheep i.d. | Sex | Source | Age | Breed | UK Meteorological<br>Season<br>Died/Euthanised | Neutered<br>Status | Staining |
| --- | --- | --- | --- | --- | --- | --- | --- |
| L44 | Male | Diagnostic PM | 7 weeks | Mule Cross | Spring | Intact | CUBIC (SMA), CUBIC (IBA1/SMA) |
| L45 | Male | Research | 3 days | Welsh<br>Mountain | Spring | Intact | CUBIC (SMA), CUBIC (IBA1/SMA) |
| S84 | Male | Research | Mature (2 years +) | Welsh<br>Mountain | Winter | Intact | Ki67/SMA, AR/SMA, ER/SMA, PR/SMA, IBA1/SMA, CD3/CD20,<br>cKit/SMA |
| S131 | Male | Diagnostic PM | 6 months | Shetland | Summer | Neutered | Ki67/SMA, AR/SMA, ER/SMA, PR/SMA, IBA1/SMA, CD3/CD20,<br>cKit/SMA |
| S132 | Male | Diagnostic PM | Mature (2 years +) | Unknown | Autumn | Intact | Ki67/SMA, AR/SMA, ER/SMA, PR/SMA, IBA1/SMA, CD3/CD20,<br>cKit/SMA |
| S133 | Male | Diagnostic PM | 9 months | Unknown | Winter | Neutered | IBA1/SMA |
| S154 | Male | Diagnostic PM | Mature (2 years +) | Unknown | Spring | Intact | Ki67/SMA, AR/SMA, ER/SMA, PR/SMA, IBA1/SMA, CD3/CD20,<br>cKit/SMA |
| P20/97 | Male | Diagnostic PM | 4 months | Mule | Summer | Neutered | Ki67/SMA, AR/SMA, ER/SMA, PR/SMA, IBA1/SMA, CD3/CD20,<br>cKit/SMA |
| P20/154 | Male | Diagnostic PM | Mature (3 years) | Suffolk | Autumn | Intact | Ki67/SMA, IBA1/SMA, cKit/SMA |
| P21/357 | Male | Diagnostic PM | 8 months | Cheviot Cross | Autumn | Neutered | Ki67/SMA, AR/SMA, ER/SMA, PR/SMA, IBA1/SMA, CD3/CD20,<br>cKit/SMA |
| P21/358 | Male | Diagnostic PM | 8 months | New Zealand<br>Romney | Autumn | Neutered | Ki67/SMA, AR/SMA, ER/SMA, PR/SMA, IBA1/SMA, CD3/CD20,<br>cKit/SMA |
| P21/366 | Male | Diagnostic PM | 9 months | Cheviot Cross | Winter | Neutered | Ki67/SMA, AR/SMA, ER/SMA, PR/SMA, IBA1/SMA, CD3/CD20,<br>cKit/SMA |
| P21/367 | Male | Diagnostic PM | 10 months | Cheviot Cross | Winter | Neutered | Ki67/SMA, AR/SMA, ER/SMA, PR/SMA, IBA1/SMA, CD3/CD20,<br>cKit/SMA |
| P21/369 | Male | Diagnostic PM | 11 months | Cheviot Cross | Winter | Neutered | Ki67/SMA, AR/SMA, ER/SMA, PR/SMA, IBA1/SMA, CD3/CD20 |
| P22/66 | Male | Diagnostic PM | 14 months | Welsh<br>Mountain | Spring | Intact | Ki67/SMA, AR/SMA, ER/SMA, PR/SMA, IBA1/SMA, CD3/CD20,<br>cKit/SMA |
| P22/27 | Male | Diagnostic PM | 8 months | Texel Cross | Winter | Neutered | Ki67/SMA, AR/SMA, ER/SMA, PR/SMA, IBA1/SMA, CD3/CD20,<br>cKit/SMA |
| P22/233 | Male | Diagnostic PM | 5 months | Texel Cross | Autumn | Intact | Ki67/SMA, AR/SMA, ER/SMA, PR/SMA, IBA1/SMA, CD3/CD20,<br>cKit/SMA |
| P23/130 | Male | Diagnostic PM | 7 weeks | Texel Cross | Spring | Neutered | CUBIC (SMA), CUBIC (IBA1/SMA) |
| P23/133 | Male | Diagnostic PM | Mature (2 years +) | Jacob Cross | Spring | Neutered | Ki67/SMA, AR/SMA, ER/SMA, PR/SMA, IBA1/SMA, CD3/CD20 |
| P16 138 | Female | Diagnostic PM | 5 months | Texel Cross | Summer | Intact | Ki67/SMA, AR/SMA, ER/SMA, PR/SMA |
| P18 247 | Female | Diagnostic PM | 4 months | Beltex | Autumn | Intact | Ki67/SMA, AR/SMA, ER/SMA, PR/SMA |
| S43 | Female | Research | 8 months | Welsh<br>Mountain | Winter | Intact | Ki67/SMA, AR/SMA, ER/SMA, PR/SMA |
| S44 | Female | Research | 11 months | Welsh<br>Mountain | Winter | Intact | Ki67/SMA, AR/SMA, ER/SMA, PR/SMA |
| S47 | Female | Research | 9 months | Welsh<br>Mountain | Winter | Intact | Ki67/SMA, AR/SMA, ER/SMA, PR/SMA |
| S52 | Female | Research | 9 months | Welsh<br>Mountain | Spring | Intact | Ki67/SMA, AR/SMA, ER/SMA, PR/SMA |
| S78 | Female | Research | 11 months | Welsh<br>Mountain | Autumn | Intact | Ki67/SMA, AR/SMA, ER/SMA, PR/SMA |
| S141 | Female | Research | Mature (2 years +) | Welsh<br>Mountain | Summer | Intact | Ki67/SMA, AR/SMA, ER/SMA, PR/SMA |
| S142 | Female | Research | Mature (2 years +) | Welsh<br>Mountain | Summer | Intact | Ki67/SMA, AR/SMA, ER/SMA, PR/SMA |
| S143 | Female | Research | Mature (2 years +) | Welsh Mountain | Summer | Intact | Ki67/SMA, AR/SMA, ER/SMA, PR/SMA |
| S144 | Female | Research | Mature (2 years +) | Welsh Mountain | Summer | Intact | Ki67/SMA, AR/SMA, ER/SMA, PR/SMA |
| S145 | Female | Research | Mature (2 years +) | Welsh Mountain | Summer | Intact | Ki67/SMA, AR/SMA, ER/SMA, PR/SMA |
| S146 | Female | Research | Mature (2 years +) | Welsh Mountain | Summer | Intact | Ki67/SMA, AR/SMA, ER/SMA, PR/SMA |
| S147 | Female | Research | Mature (2 years +) | Welsh Mountain | Summer | Intact | Ki67/SMA, AR/SMA, ER/SMA, PR/SMA |

**Supplementary Table 2:** Full details of primary and secondary antibodies utilised in immunohistochemical, immunofluorescence and CUBIC staining.

| Target | Application (IHC, immunohistochemistry; IF, immunofluorescence; CUBIC, 3D tissue clearing) | Species and Clone (where stated) | Dilution | Manufacturer | Catalogue number |
| --- | --- | --- | --- | --- | --- |
| *Primary antibodies* |  |  |  |  |  |
| E-cadherin | IF | Mouse monoclonal anti-human [NCH-38] | 1:100 | Dako/Agilent | M3612 |
| Alpha Smooth muscle actin | Dual colour IHC, CUBIC | Mouse monoclonal anti-human [1A4] | 1:500 (dual colour IHC), 1:100 (CUBIC) | Dako/Agilent | M0851 |
| Alpha Smooth muscle actin | Dual colour IHC, IF | Rabbit monoclonal [EPR5368] | 1:2000 | Abcam | Ab124964 |
| Ki67 | Dual colour IHC | Mouse monoclonal anti-human [MIB-1] | 1:100 | Dako/Agilent | M7240 |
| Androgen Receptor | IF | Rabbit monoclonal [EPR1535(2)] | 1:100 | Abcam | Ab133273 |
| Oestrogen Receptor $\alpha$ | IF | Rabbit monoclonal [D6R2W] | 1:200 | Cell Signalling Technology | 13258S |
| Progesterone Receptor A/B | IF | Rabbit monoclonal [D8Q2J] | 1:1000 | Cell Signalling Technology | 8757S |
| IBA1 | Dual colour IHC, CUBIC | Rabbit monoclonal [EPR16588] | 1:1200 (dual colour IHC), 1:400 (CUBIC) | Abcam | Ab178846 |
| CD3 | Dual colour IHC | Mouse monoclonal [F7.2.38] | 1:250 | Dako/Agilent | M7254 |
| CD20 | Dual colour IHC | Rabbit monoclonal [E7B7T] | 1:500 | Cell Signalling Technology | 48750S |
| c-Kit | Dual colour IHC | Rabbit monoclonal [D3W6Y] | 1:100 | Cell Signalling Technology | 37805S |
| *Secondary antibodies* |  |  |  |  |  |
| Mouse IgG, Alexa Fluor Plus 488 | IF, CUBIC | Goat | 1:500 | Thermo Fisher Scientific | A32723 |
| Rabbit IgG, Alexa Fluor Plus 647 | IF, CUBIC | Goat | 1:500 | Thermo Fisher Scientific | A32733 |

## Notes

### Competing Interest Statement

The authors have declared no competing interest.

## References

Atwood, C.S., Hovey, R.C., Glover, J.P., Chepko, G., Ginsburg, E., Robison, W.G., Vonderhaar, B.K., 2000. Progesterone induces side-branching of the ductal epithelium in the mammary glands of peripubertal mice. Journal of Endocrinology 167, 39–52. 10.1677/joe.0.1670039

Beaudry, K.L., Parsons, C.L.M., Ellis, S.E., Akers, R.M., 2016. Localization and quantitation of macrophages, mast cells, and eosinophils in the developing bovine mammary gland1. Journal of Dairy Science 99, 796–804. 10.3168/jds.2015-9972

Brisken, C., Park, S., Vass, T., Lydon, J.P., O’Malley, B.W., Weinberg, R.A., 1998. A paracrine role for the epithelial progesterone receptor in mammary gland development. Proc Natl Acad Sci U S A 95, 5076–5081.

Cardiff, R.D., Jindal, S., Treuting, P.M., Going, J.J., Gusterson, B., Thompson, H.J., 2018. 23

-Mammary Gland, in: Treuting, P.M., Dintzis, S.M., Montine, K.S. (Eds.), Comparative Anatomy and Histology (Second Edition). Academic Press, San Diego, pp. 487–509. 10.1016/B978-0-12-802900-8.00023-3

Cardoso, F., Bartlett, J.M.S., Slaets, L., van Deurzen, C.H.M., van Leeuwen-Stok, E., Porter, P., Linderholm, B., Hedenfalk, I., Schröder, C., Martens, J., Bayani, J., van Asperen, C., Murray, M., Hudis, C., Middleton, L., Vermeij, J., Punie, K., Fraser, J., Nowaczyk, M., Rubio, I.T., Aebi, S., Kelly, C., Ruddy, K.J., Winer, E., Nilsson, C., Lago, L.D., Korde, L., Benstead, K., Bogler, O., Goulioti, T., Peric, A., Litière, S., Aalders, K.C., Poncet, C., Tryfonidis, K., Giordano, S.H., 2018. Characterization of male breast cancer: results of the EORTC 10085/TBCRC/BIG/NABCG International Male Breast Cancer Program. Annals of Oncology, Incorporating blood-based liquid biopsy information into cancer staging 29, 405–417. 10.1093/annonc/mdx651

Chatterji, S., Krzoska, E., Thoroughgood, C.W., Saganty, J., Liu, P., Elsberger, B., Abu-Eid, R., Speirs, V., 2023. Defining genomic, transcriptomic, proteomic, epigenetic, and phenotypic biomarkers with prognostic capability in male breast cancer: a systematic review. The Lancet Oncology 24, e74–e85. 10.1016/S1470-2045(22)00633-7

Chen, W., Beck, I., Schober, W., Brockow, K., Effner, R., Buters, J.T.M., Behrendt, H., Ring, J., 2010. Human mast cells express androgen receptors but treatment with testosterone exerts no influence on IgE-independent mast cell degranulation elicited by neuromuscular blocking agents. Experimental Dermatology 19, 302–304. 10.1111/j.1600-0625.2009.00969.x

Chua, A.C.L., Hodson, L.J., Moldenhauer, L.M., Robertson, S.A., Ingman, W.V., 2010. Dual roles for macrophages in ovarian cycle-associated development and remodelling of the mammary gland epithelium. Development 137, 4229–4238. 10.1242/dev.059261

Colitti, M., Parillo, F., 2013. Immunolocalization of estrogen and progesterone receptors in ewe mammary glands. Microscopy Research and Technique 76, 955–962. 10.1002/jemt.22254

Dawson, C.A., Pal, B., Vaillant, F., Gandolfo, L.C., Liu, Z., Bleriot, C., Ginhoux, F., Smyth, G.K., Lindeman, G.J., Mueller, S.N., Rios, A.C., Visvader, J.E., 2020. Tissue-resident ductal macrophages survey the mammary epithelium and facilitate tissue remodelling. Nat Cell Biol 22, 546–558. 10.1038/s41556-020-0505-0

Dimitrakakis, C., Bondy, C., 2009. Androgens and the breast. Breast Cancer Research 11, 212. 10.1186/bcr2413

Drews, Ulrich, Drews, Ute, 1977. Regression of mouse mammary gland anlagen in recombinants of Tfm and wild-type tissues: testosterone acts via the mesenchyme. Cell 10, 401–404. 10.1016/0092-8674(77)90027-7

Dürnberger, H., Kratochwil, K., 1980. Specificity of tissue interaction and origin of mesenchymal cells in the androgen response of the embryonic mammary gland. Cell 19, 465–471. 10.1016/0092-8674(80)90521-8

Fox, S., Speirs, V., Shaaban, A.M., 2022. Male breast cancer: an update. Virchows Arch 480, 85–93. 10.1007/s00428-021-03190-7

Gao, Y.R. (Ellen), Walters, K.A., Desai, R., Zhou, H., Handelsman, D.J., Simanainen, U., 2014. Androgen Receptor Inactivation Resulted in Acceleration in Pubertal Mammary Gland Growth, Upregulation of ERα Expression, and Wnt/β-Catenin Signaling in Female Mice. Endocrinology 155, 4951–4963. 10.1210/en.2014-1226

Hassiotou, F., Geddes, D., 2013. Anatomy of the human mammary gland: Current status of knowledge. Clin. Anat. 26, 29–48. 10.1002/ca.22165

Hilton, H.N., Graham, J.D., Clarke, C.L., 2015. Minireview: Progesterone Regulation of Proliferation in the Normal Human Breast and in Breast Cancer: A Tale of Two Scenarios? Molecular Endocrinology 29, 1230–1242. 10.1210/me.2015-1152

Hodson, L.J., Chua, A.C.L., Evdokiou, A., Robertson, S.A., Ingman, W.V., 2013. Macrophage Phenotype in the Mammary Gland Fluctuates over the Course of the Estrous Cycle and Is Regulated by Ovarian Steroid Hormones1. Biology of Reproduction 89, 65, 1–8. 10.1095/biolreprod.113.109561

Hovey, R.C., Mcfadden, T.B., Akers, R.M., 1999. Regulation of Mammary Gland Growth and Morphogenesis by the Mammary Fat Pad: A Species Comparison. J Mammary Gland Biol Neoplasia 4, 53–68. 10.1023/A:1018704603426

Hughes, K., 2023. Studying Mammary Physiology and Pathology in Domestic Species Benefits Both Humans and Animals. J Mammary Gland Biol Neoplasia 28, 18. 10.1007/s10911-023-09547-9

Hughes, K., 2021a. Development and Pathology of the Equine Mammary Gland. J Mammary Gland Biol Neoplasia 26, 121–134. 10.1007/s10911-020-09471-2

Hughes, K., 2021b. Comparative mammary gland postnatal development and tumourigenesis in the sheep, cow, cat and rabbit: Exploring the menagerie. Semin Cell Dev Biol 114, 186–195. 10.1016/j.semcdb.2020.09.010

Hughes, K., Watson, C.J., 2018. The Mammary Microenvironment in Mastitis in Humans, Dairy Ruminants, Rabbits and Rodents: A One Health Focus. J Mammary Gland Biol Neoplasia 23, 27–41. 10.1007/s10911-018-9395-1

Hughes, K., Wickenden, J.A., Allen, J.E., Watson, C.J., 2012. Conditional deletion of Stat3 in mammary epithelium impairs the acute phase response and modulates immune cell numbers during post-lactational regression. The Journal of Pathology 227, 106–117. 10.1002/path.3961

Inman, J.L., Robertson, C., Mott, J.D., Bissell, M.J., 2015. Mammary gland development: cell fate specification, stem cells and the microenvironment. Development 142, 1028–1042. 10.1242/dev.087643

Iuanow, E., Kettler, M., Slanetz, P.J., 2011. Spectrum of Disease in the Male Breast. American Journal of Roentgenology 196, W247–W259. 10.2214/AJR.09.3994

Jenkinson, C.M.C., 2003. The pattern and regulation of mammary gland development during fetal life in sheep : a thesis presented in partial fulfilment of the requirements for the degree of Doctor of Philosophy in Animal Science at Massey University, Palmerston North, New Zealand. Massey University.

Jesinger, R.A., 2014. Breast Anatomy for the Interventionalist. Techniques in Vascular and Interventional Radiology, Breast Interventions 17, 3–9. 10.1053/j.tvir.2013.12.002

Kolla, S., Pokharel, A., Vandenberg, L.N., 2017. The mouse mammary gland as a sentinel organ: distinguishing ‘control’ populations with diverse environmental histories. Environ Health 16, 25. 10.1186/s12940-017-0229-1

Lammie, A., Drobnjak, M., Gerald, W., Saad, A., Cote, R., Cordon-Cardo, C., 1994. Expression of c-kit and kit ligand proteins in normal human tissues. Journal of Histochemistry & Cytochemistry 42, 1417–1425. 10.1177/42.11.7523489

Lilla, J.N., Werb, Z., 2010. Mast cells contribute to the stromal microenvironment in mammary gland branching morphogenesis. Developmental Biology 337, 124–133. 10.1016/j.ydbio.2009.10.021

Liu, N., Johnson, K.J., Ma, C.X., 2018. Male Breast Cancer: An Updated Surveillance, Epidemiology, and End Results Data Analysis. Clinical Breast Cancer 18, e997–e1002. 10.1016/j.clbc.2018.06.013

Liva, S.M., Voskuhl, R.R., 2001. Testosterone Acts Directly on CD4+ T Lymphocytes to Increase IL-10 Production1. The Journal of Immunology 167, 2060–2067. 10.4049/jimmunol.167.4.2060

Lloyd-Lewis, B., Davis, F.M., Harris, O.B., Hitchcock, J.R., Lourenco, F.C., Pasche, M., Watson, C.J., 2016. Imaging the mammary gland and mammary tumours in 3D: optical tissue clearing and immunofluorescence methods. Breast Cancer Research 18, 127. 10.1186/s13058-016-0754-9

Macias, H., Hinck, L., 2012. Mammary gland development. WIREs Developmental Biology 1, 533–557. 10.1002/wdev.35

Morrison, J., Berry, M., Botting, K., Darby, J., Frasch, M., Gatford, K., Giussani, D., Gray, C., Harding, R., Herrera, E., Kemp, M., Lock, M., Mcmillen, I., Moss, T., Musk, G., Oliver, M., Regnault, T., Roberts, C., Soo, J., Tellam, R., 2018. Improving pregnancy outcomes in humans through studies in sheep. American Journal of Physiology-Regulatory, Integrative and Comparative Physiology 315. 10.1152/ajpregu.00391.2017

Nagy, D., Gillis, C.M.C., Davies, K., Fowden, A.L., Rees, P., Wills, J.W., Hughes, K., 2021. Developing ovine mammary terminal duct lobular units have a dynamic mucosal and stromal immune microenvironment. Commun Biol 4, 993. 10.1038/s42003-021-02502-6

Newman, S.J., Smith, S.A., Zimmerman, K., 2021. Mammary carcinoma arising in an adenoma in a ewe. J Vet Diagn Invest 33, 566–571. 10.1177/1040638721993061

Paine, I.S., Lewis, M.T., 2017. The Terminal End Bud: the Little Engine that Could. J Mammary Gland Biol Neoplasia 22, 93–108. 10.1007/s10911-017-9372-0

Pokharel, A., Kolla, S., Matouskova, K., Vandenberg, L.N., 2018. Asymmetric development of the male mouse mammary gland and its response to a prenatal or postnatal estrogen challenge. Reprod Toxicol 82, 63–71. 10.1016/j.reprotox.2018.10.003

Ramirez, R.A., Lee, A., Schedin, P., Russell, J.S., Masso-Welch, P.A., 2012. Alterations in mast cell frequency and relationship to angiogenesis in the rat mammary gland during windows of physiologic tissue remodeling. Developmental Dynamics 241, 890–900. 10.1002/dvdy.23778

Renfree, M.B., Robinson, E.S., Short, R.V., Vandeberg, J.L., 1990. Mammary glands in male marsupials: I. Primordia in neonatal opossums Didelphis virginiana and Monodelphis domestica. Development 110, 385–390. 10.1242/dev.110.2.385

Ribatti, D., 2018. The Staining of Mast Cells: A Historical Overview. International Archives of Allergy and Immunology 176, 55–60. 10.1159/000487538

Richert, M.M., Schwertfeger, K.L., Ryder, J.W., Anderson, S.M., 2000. An Atlas of Mouse Mammary Gland Development. J Mammary Gland Biol Neoplasia 5, 227–241. 10.1023/A:1026499523505

Rowson, A.R., Daniels, K.M., Ellis, S.E., Hovey, R.C., 2012. Growth and development of the mammary glands of livestock: A veritable barnyard of opportunities. Seminars in Cell & Developmental Biology, Cell Regulation by Selective Protein Degradation & Biology of Mammary Gland Development 23, 557–566. 10.1016/j.semcdb.2012.03.018

Safayi, S., Korn, N., Bertram, A., Akers, R.M., Capuco, A.V., Pratt, S.L., Ellis, S., 2012. Myoepithelial cell differentiation markers in prepubertal bovine mammary gland: Effect of ovariectomy. Journal of Dairy Science 95, 2965–2976. 10.3168/jds.2011-4690

Schindelin, J., Arganda-Carreras, I., Frise, E., Kaynig, V., Longair, M., Pietzsch, T., Preibisch, S., Rueden, C., Saalfeld, S., Schmid, B., Tinevez, J.-Y., White, D.J., Hartenstein, V., Eliceiri, K., Tomancak, P., Cardona, A., 2012. Fiji: an open-source platform for biological-image analysis. Nature Methods 9, 676–682. 10.1038/nmeth.2019

Simanainen, U., Gao, Y.R., Walters, K.A., Watson, G., Desai, R., Jimenez, M., Handelsman, D.J., 2012. Androgen Resistance in Female Mice Increases Susceptibility to DMBA-Induced Mammary Tumors. Hormones and Cancer 3, 113–124. 10.1007/s12672-012-0107-9

Stewart, T.A., Hughes, K., Hume, D.A., Davis, F.M., 2019. Developmental Stage-Specific Distribution of Macrophages in Mouse Mammary Gland. Frontiers in Cell and Developmental Biology 7, 250. 10.3389/fcell.2019.00250

Susaki, E.A., Tainaka, K., Perrin, D., Kishino, F., Tawara, T., Watanabe, T.M., Yokoyama, C., Onoe, H., Eguchi, M., Yamaguchi, S., Abe, T., Kiyonari, H., Shimizu, Y., Miyawaki, A., Yokota, H., Ueda, H.R., 2014. Whole-Brain Imaging with Single-Cell Resolution Using Chemical Cocktails and Computational Analysis. Cell 157, 726–739. 10.1016/j.cell.2014.03.042

Szabo, G.K., Vandenberg, L.N., 2021. REPRODUCTIVE TOXICOLOGY: The male mammary gland: a novel target of endocrine-disrupting chemicals. Reproduction 162, F79– F89. 10.1530/REP-20-0615

Tower, H., Dall, G., Davey, A., Stewart, M., Lanteri, P., Ruppert, M., Lambouras, M., Nasir, I., Yeow, S., Darcy, P.K., Ingman, W.V., Parker, B., Haynes, N.M., Britt, K.L., 2022. Estrogen-induced immune changes within the normal mammary gland. Scientific Reports 12, 18986. 10.1038/s41598-022-21871-4

Vandenberg, L.N., Schaeberle, C.M., Rubin, B.S., Sonnenschein, C., Soto, A.M., 2013. The male mammary gland: A target for the xenoestrogen bisphenol A. Reproductive Toxicology 37, 15–23. 10.1016/j.reprotox.2013.01.002

